# A role for the putative error-prone polymerase *REV1* in DNA damage and antifungal drug resistance in *Candida albicans*

**DOI:** 10.1101/2024.06.24.600412

**Authors:** Michelle R. Agyare-Tabbi, Deeva Uthayakumar, Desiree Francis, Laetitia Maroc, Chris Grant, Peter McQueen, Garret Westmacott, Hajer Shaker, Iwona Skulska, Isabelle Gagnon-Arsenault, Jonathan Boisvert, Christian R. Landry, Rebecca S. Shapiro

**Affiliations:** Department of Molecular and Cellular Biology, University of Guelph, Guelph, Ontario, Canada N1H 5N4; Department of Immunology, University of Toronto, Toronto, Ontario, Canada M5R 0A3; National Microbiology Laboratory, Public Health Agency of Canada, Winnipeg, Manitoba, Canada, R3E 3L5; Institut de Biologie Intégrative et des Systèmes (IBIS), Université Laval, Québec, QC, Canada; Department of Biochemistry, Microbiology and Bioinformatics, Université Laval, Québec, QC, Canada; Quebec Network for Research on Protein Function, Engineering, and Applications (PROTEO), Université du Québec à Montréal, Montréal, QC, Canada; Université Laval Big Data Research Center (BDRC_UL), Québec, QC, Canada

## Abstract

Antimicrobial-induced DNA damage, and subsequent repair via upregulation of DNA repair factors, including error-prone translesion polymerases, can lead to the increased accumulation of mutations in the microbial genome, and ultimately increased risk of acquired mutations associated with antimicrobial resistance. While this phenotype is well described in bacterial species, it is less thoroughly investigated amongst microbial fungi. Here, we monitor DNA damage induced by antifungal agents in the fungal pathogen *Candida albicans*, and find that commonly used antifungal drugs are able to induce DNA damage, leading to the upregulation of transcripts encoding predicted error-prone polymerases and related factors. We focus on *REV1*, encoding a putative error-prone polymerase, and find that while deleting this gene in *C. albicans* leads to increased sensitivity to DNA damage, it also unexpectedly renders cells more likely to incur mutations and evolve resistance to antifungal agents. We further find that deletion of *REV1* leads to a significant depletion in the uncharacterized protein Shm1, which itself plays a role in fungal mutagenesis. Together, this work lends new insight into previously uncharacterized factors with important roles in the DNA damage response, mutagenesis, and the evolution of antifungal drug resistance.

## Introduction

Fungal infections pose a major threat to human health and are responsible for 150 million life-threatening infections and 1.5 million annual deaths^1^. Fungi are ubiquitous pathogens, responsible for a broad range of diseases, from common skin and mucosal infections to life-threatening invasive mycoses, which pose a distinct threat among people with underlying conditions or immunodeficiencies, including transplant and cancer patients. Global rates of fungal infections have been increasing as people belonging to vulnerable risk groups increase, and as climate change continues to influence the spread and emergence of fungal pathogens^1–5^. Amongst these fungal pathogens, *Candida* species are the leading cause of invasive infections^6^ and the fourth most common cause of hospital-acquired bloodstream infections^7^. The opportunistic pathogen *Candida albicans* is the predominant cause of candidiasis diseases, although non-*albicans Candida* pathogens represent a growing proportion of clinically isolated species^8–10^, which are characterized by high rates of antifungal drug resistance^9,11^.

Treatment for fungal infections, including *C. albicans* infections, is difficult due to an overall paucity of effective antifungal therapies, despite the growing importance of fungal infections. A major challenge in identifying novel antifungal therapies is the occurrence of off-target effects and host toxicity due to the conserved eukaryotic biology of human hosts and fungi^12^. In addition to this, antifungal drug resistance has been identified as a major threat to human health, as *C. albicans* and other *Candida* pathogens are increasingly identified with reduced susceptibility to antifungal agents^11^. Currently, three major classes of antifungal therapies are clinically available for the treatment of *C. albicans* infections: polyenes, azoles, and echinocandins^12^. The most recently discovered class of antifungal drugs is the echinocandins, considered the first-line therapy for invasive candidiasis^13^. Echinocandins exhibit fungicidal activity against *Candida* pathogens and target the fungal cell wall by inhibiting ß-(1,3)-D-glucan synthase (encoded by *FKS1/2*), an enzyme critical to cell wall synthesis^14^. In this class, the most commonly used drug against *Candida* infections is caspofungin^14^. Resistance to echinocandin antifungals has been increasingly found amongst fungal clinical isolates, particularly *Nakaseomyces* (*Candida*) *glabrata* and *Candida auris* strains^15^. The limited number of antifungal therapies, and the rising global threat that fungal pathogens pose to human health, highlight a need for robust characterization of antifungal drug resistance evolution in fungal pathogens.

For bacterial species, treatment with antimicrobial agents has been shown to play a role in DNA damage, mutagenesis, and the subsequent development of antibiotic drug resistance^16–20^. Antibiotics can induce DNA damage in bacteria either directly, as is the case with fluoroquinolones that cause DNA damage by inhibiting DNA topoisomerases^21,22^, or more indirectly by promoting the production of cellular reactive oxygen species (ROS), which in turn cause damage to DNA^23–25^. Various cellular stress responses become upregulated upon treatment with such antibiotics, including the DNA damage-induced SOS response. The bacterial SOS response induces the expression of error-prone DNA translesion polymerases to manage the DNA damage^16,18,19,26–30^. These polymerases have reduced fidelity compared to replicative polymerases, therefore accelerating the introduction of mutations into the genome, which in turn provides more mutational variation that may confer resistance to antibiotics^16,18,19,29,30^. In fungal pathogens, there is evidence that similar to antibiotics, antifungal agents are capable of inducing ROS production and DNA damage^31–33^. Further, in *C. albicans*, antifungal agents can increase the rate of genetic alterations in fungi, including point mutations^34^ and loss of heterozygosity (LOH)^35^.

Despite evidence for antifungal-induced DNA damage and mutagenesis in fungi, the role of error-prone polymerases in this process has not been investigated in fungal pathogens. However, in the model yeast *Saccharomyces cerevisiae*, the role of error-prone polymerases has been well described^36^. In *S. cerevisiae*, translesion synthesis (TLS) is mediated by three main error-prone TLS polymerases: Rev1, Polζ (consisting of Rev3 and Rev7 subunits), and Polη (Rad30)^37^. These TLS polymerases are broadly characterized by their ability to replicate damaged DNA via TLS, whereby a DNA lesion is bypassed by the incorporation of a nucleotide opposite the lesion^37^. While many DNA lesions cannot be used as a template by stringent replicative polymerases, TLS polymerases are able to use damaged DNA as a template ^37^. Compared with replicative polymerases, which exploit rigorous proofreading activity and have extremely low error rates (∼1 incorrect nucleotide per 10^6^-10^8^ bases), the error-prone TLS polymerases lack 3’-to-5’ proofreading activity and have very high error rates (∼1 incorrect nucleotide per 10^1^-10^4^ bases)^37,38^. Rev1 and Polζ often act in concert^39,40^ and in *S. cerevisiae*, the Rev1/Polζ complex is a central regulator of translesion synthesis, introducing errors in both damaged and undamaged DNA, and is considered to be responsible for a majority of all mutations in this yeast^39,41,42^. Deletion of *REV1* in *S. cerevisiae* renders cells highly susceptible to DNA damage and decreases the rate at which mutations are acquired^43–45^. In *C. albicans*, however, the role of Rev1 and other error-prone TLS polymerases has not been closely investigated.

Here, we characterize the role of the putative TLS polymerase Rev1 in the fungal pathogen *C. albicans*, and investigate the role of Rev1 in mutagenesis. We confirm and quantify *C. albicans* DNA damage in the presence of antifungal drugs, and further find that numerous predicted error-prone polymerases (and related factors) are upregulated in the presence of DNA-damaging agents, including antifungal drugs. We further focus on *REV1*, which when deleted, leads to an increased mutation rate (in opposition to what is observed in *S. cerevisiae*^43–45^), and a corresponding increase in evolved resistance to the antifungal drug caspofungin. Deletion of *REV1* also causes a higher abundance of the protein Shm1, which is also involved in the *C. albicans* mutation rate. Together this work characterizes new factors involved in DNA damage and mutagenesis in *C. albicans*.

## Results

### Antifungal drugs induce DNA double-strand breaks in C. albicans

To assess the role of TLS polymerases on antifungal-induced mutagenesis in *C. albicans*, we first sought to assess and quantify DNA damage induced in fungal cells upon treatment with antifungal agents. For this, we exploited a *C. albicans* strain that we previously generated containing a *GAM-GFP* construct to visualize and quantify DNA damage^46^. Gam is a viral protein that binds irreversibly to DNA double-strand breaks (DSBs), and fluorescent labelling of Gam can effectively visualize and quantify DNA damage in the form of DSBs in live microbial cells, including *C. albicans*^46,47^. The Gam-GFP system we used here is under the control of the *tetON* tetracycline-inducible promoter, such that the presence of tetracycline or its derivative doxycycline, leads to the production of the Gam-GFP protein.

We treated *GAM-GFP* expressing *C. albicans* cells with antifungals fluconazole or caspofungin and included treatment with hydrogen peroxide (H_2_O_2_) as a control for induction of DNA DSBs. We co-stained cells with the nuclear stain DAPI and used co-localization of GFP and DAPI to quantify DSBs. We found a significant increase in GFP co-localization with the DAPI-stained nuclei in the cells treated with each antifungal, as well as H_2_O_2_, compared to the untreated cells (**FIGURE 1**), indicating that both azole and echinocandin antifungals are capable of inducing DNA damage in the form of DSBs in *C. albicans*.

**Figure 1.**
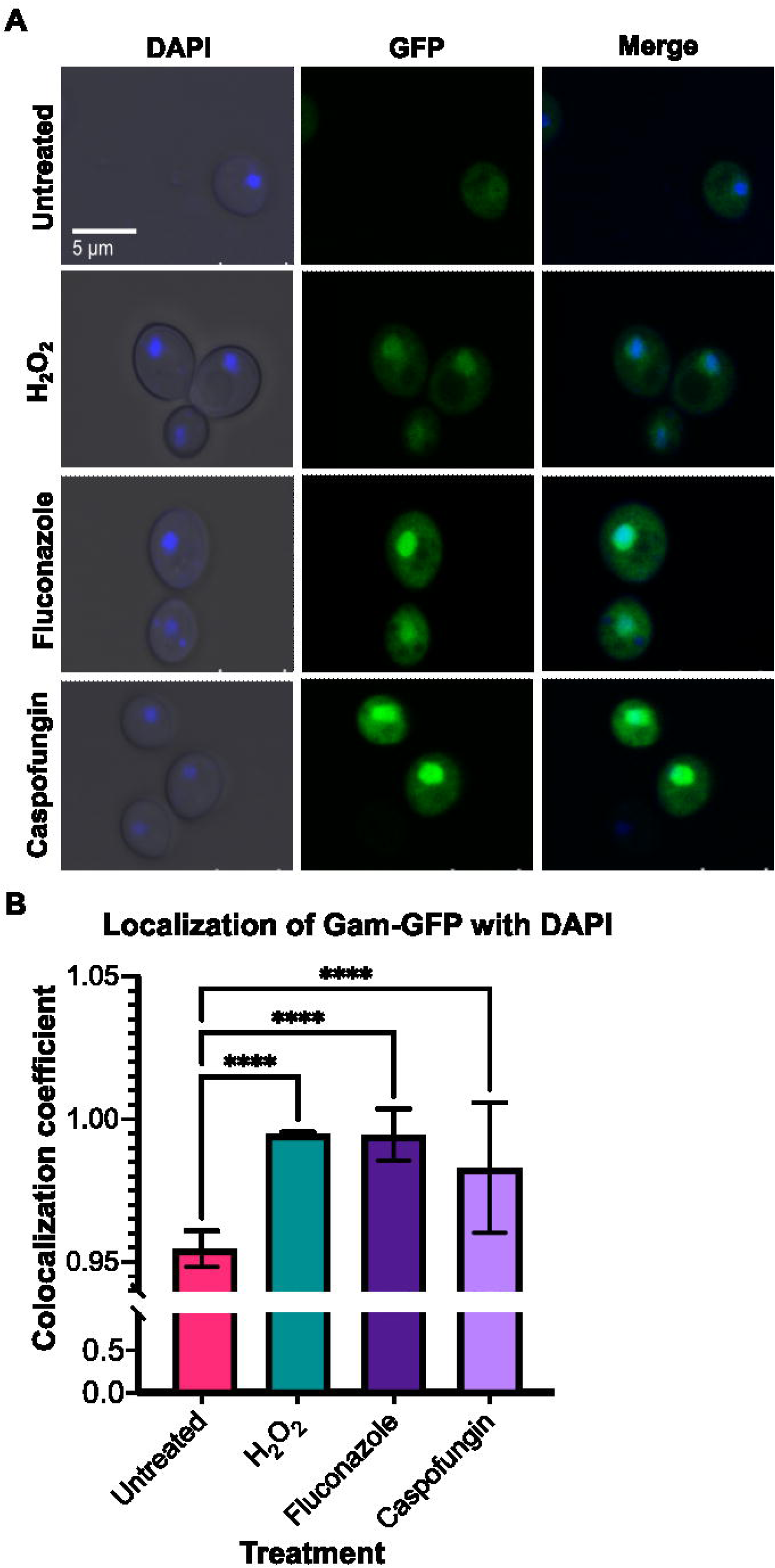
Treatment with antifungal drugs induces DNA damage. **A.** Fluorescent microscopy depicts DAPI-stained nuclei, *GAM-GFP* signal, co-localization of DAPI and GFP, and bright field (BF) images. **B.** Quantification of fluorescence microscopy displayed as a colocalization coefficient. A two-way ANOVA test was conducted and error bars represent SEM. *P* < 0.0001 (****).

### Genes encoding C. albicans TLS polymerases are upregulated in response to antifungal drugs and DNA-damaging agents

Following confirmation that treatment with antifungal drugs results in DNA DSBs, we next sought to determine whether *C. albicans* genes encoding putative TLS polymerases (and related factors) were upregulated under these stress treatment conditions. We used reverse transcriptase quantitative PCR (RT-qPCR) to measure gene expression in *C. albicans* that had been exposed to the antifungal caspofungin or methyl methanesulfonate (MMS) as a DNA-damaging alkylating agent. We monitored the expression of six target genes, chosen based on known orthologs of TLS polymerases and associated factors in *S. cerevisiae*: *REV1, REV3, RAD32, RAD5, RAD6,* and *REV7*. Upon treatment with MMS, most of the tested genes showed an increase in expression with increasing concentrations of MMS, and all genes showed significant increases in expression at the highest concentration compared with the untreated control (**FIGURE 2A**). Upon treatment with the caspofungin, *REV1, REV3,* and *RAD5* were increased in expression relative to the untreated control, with *REV1* and *RAD5* having the highest fold change in this condition (**FIGURE 2B**). Together, this suggests that several putative *C. albicans* TLS polymerases and associated factors are upregulated in the presence of DNA-damaging agents, including MMS and the antifungal caspofungin, and further reinforces our finding of caspofungin as a DNA-damaging agent.

**Figure 2.**
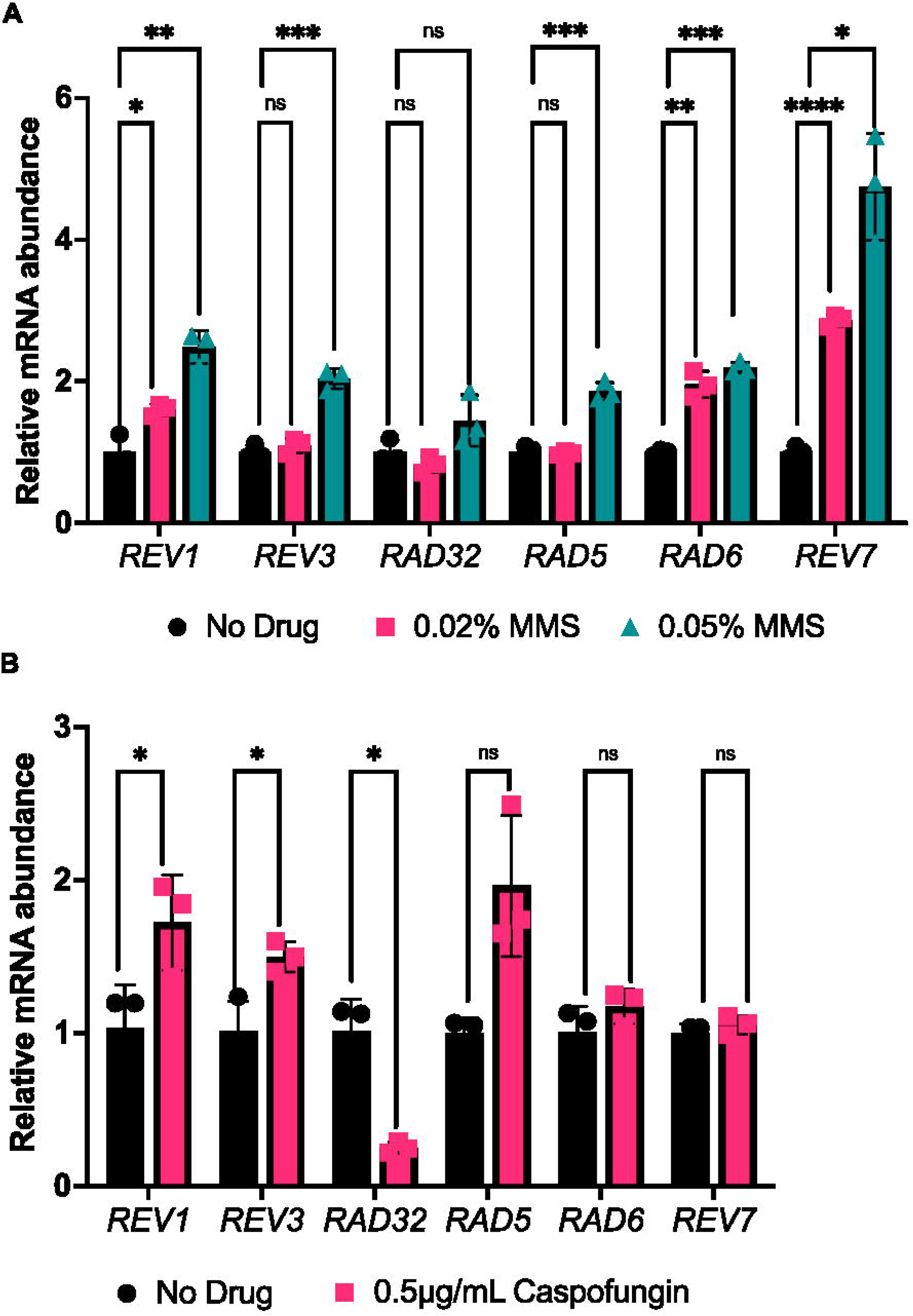
Genes encoding error-prone polymerases are upregulated when exposed to caspofungin and MMS. **A**. RT-qCR results for putative TLS polymerases in response to MMS treatment. Error bars represent SEM. **B**. RT-qCR results for putative TLS polymerases in response to caspofungin treatment. An unpaired t-test was conducted. *P* values for the significant pairwise comparisons are as follows: *P* < 0.05 (*), *P* < 0.01 (**), *P* < 0.001 (***), *P* < 0.0001 (****).

### REV1 is required for tolerance to DNA-damaging agents

Based on this observed upregulation of *REV1* in DNA-damaging conditions in *C. albicans*, and the known importance of *REV1* in *S. cerevisiae* DNA damage response and TLS^43–45^, we chose to perform a follow-up analysis on *REV1* specifically. First, we sought to determine whether *REV1* played a role in *C. albicans*’ DNA damage response. We generated a mutant strain deleted for the *REV1* gene using a CRISPR deletion strategy^48^ and monitored its sensitivity to the DNA-damaging agent MMS compared to a wild-type control strain. The rev1Δ/Δ homozygous deletion strain grew similar to the wild-type strain on rich media (**FIGURE 3**), but was severely impaired in growth in the presence of MMS (**FIGURE 3**), suggesting that similar to *S. cerevisiae*, *REV1* is essential for tolerance to DNA damage. To further confirm this observed phenotype, we knocked out *REV1* in a different *C. albicans* strain background and reconstituted the wild-type *REV1* gene back into this mutant strain. We monitored growth on rich media and MMS for both the wild-type, rev1Δ/Δ deletion, and rev1Δ/Δ::REV1 reconstituted strains, and confirmed MMS sensitivity upon deletion of *REV1*, which was restored upon reintroduction of the *REV1* gene (**FIGURE S1**).

**Figure 3.**
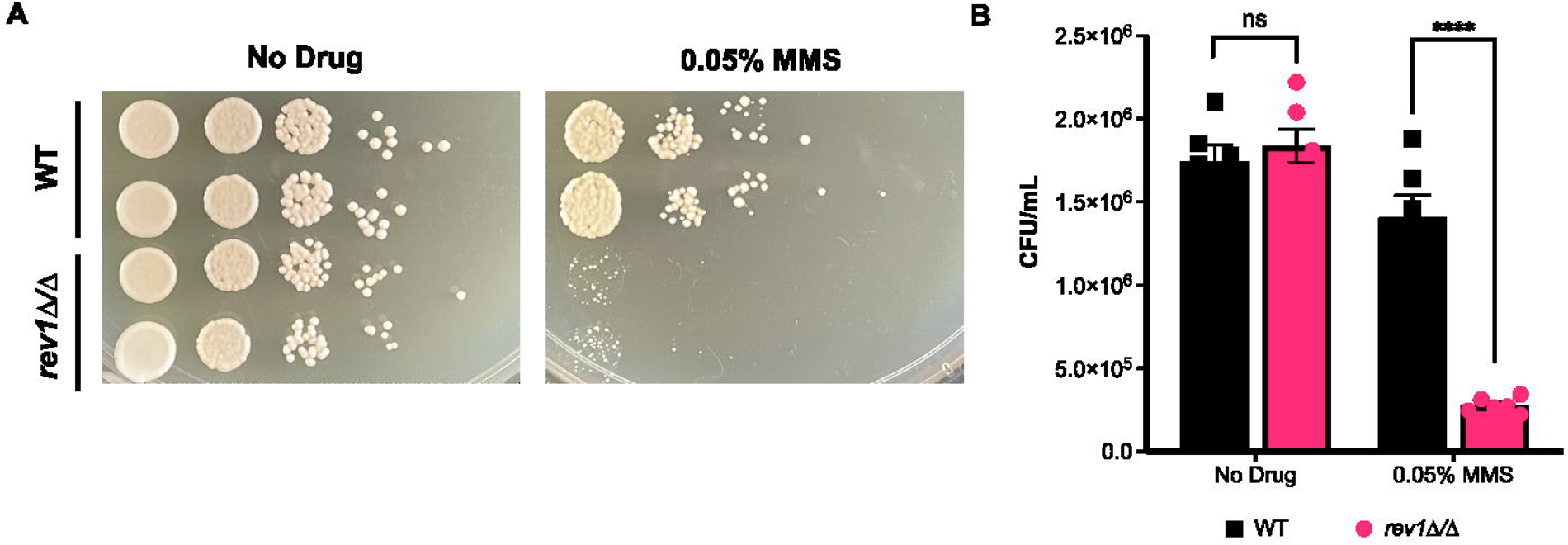
Deletion of *REV1* results in sensitivity to MMS. **A**. *REV1* deletion mutant and wild-type strains were grown in the presence of MMS and spotted on YPD. **B.** Quantification of MMS spot plates. An unpaired t-test was conducted. *P* < 0.0001 (****).

### Deletion of REV1 increases the mutation rate in C. albicans

Following confirmation of *REV1*’s role in the response to DNA damage, we next sought to determine its role in *C. albicans* mutation frequency, since work in *S. cerevisiae* indicates that deletion of *REV1* leads to a decreased mutation rate^43–45^. To assess the rate at which genetic alterations are occurring in *C. albicans*, we used a 5-fluoroorotic acid (5-FOA) fluctuation test. The 5-FOA assay is a well-established protocol^49^ used to estimate mutation rate based on the frequency at which the *URA3* gene becomes mutated and inactivated, allowing cells to grow in the presence of 5-FOA, as *URA3* encodes the enzyme for decarboxylation of 5-FOA to 5-fluorouracil, which is a toxic metabolite^49^. For this assay, we tested the rev1Δ/Δ mutant strain and a wild-type control strain, both constructed in a heterozygous URA3/ura3Δ background, to facilitate the ability to detect inactivating mutations in a single *URA3* allele. When these strains were grown and plated on 5-FOA media, the rev1Δ/Δ mutant strain produced significantly more 5-FOA resistant colonies, indicating a relatively higher rate of *URA3* mutation (**FIGURE 4**). We further confirmed this finding in the rev1Δ/Δ mutant and corresponding *REV1* constituted strain (**FIGURE S2**). This suggests that mutation of *URA3* is occurring at a higher frequency in the absence of *REV1*, which is contrary to what is seen in *S. cerevisiae*, where deletion of *REV1* leads to a decrease in *URA3* mutation rate^43–45,50^.

**Figure 4.**
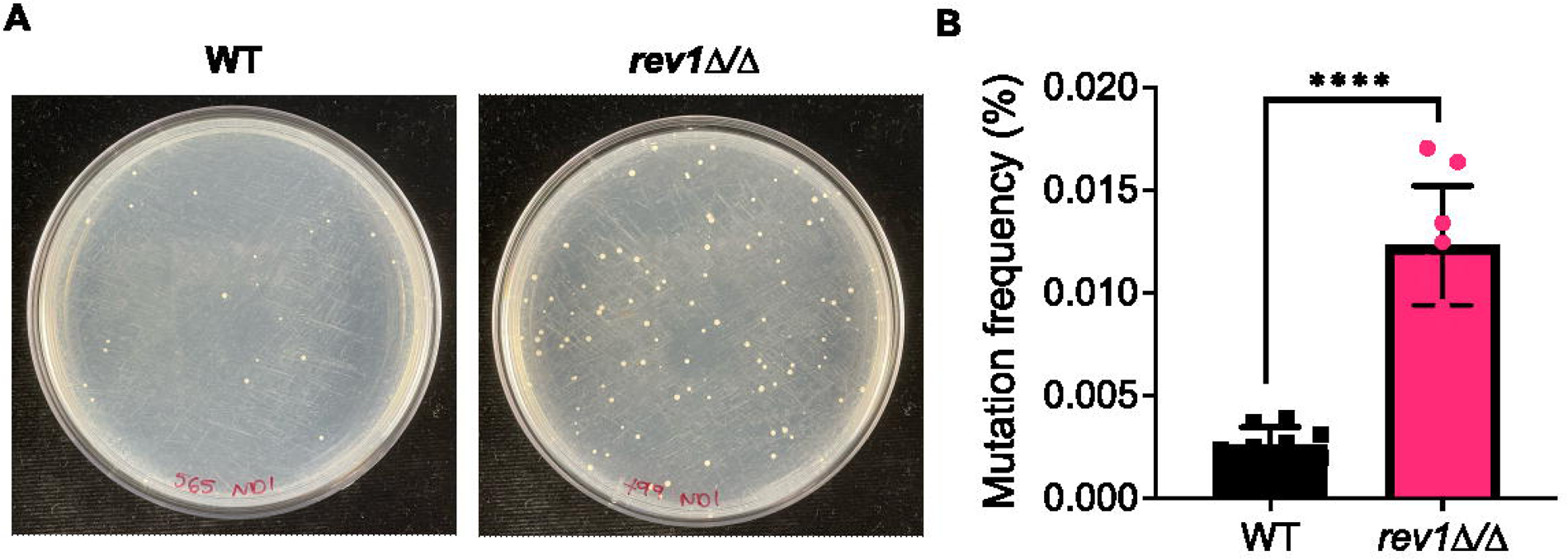
Deletion of *REV1* causes an increased frequency of mutagenesis. **A.** Plate images depict wild-type and *rev1*Δ mutant strains on 5-FOA agar. **B.** Quantified mutation rate based on the number of 5-FOA colonies as a fraction of total number of cells plated. An unpaired t-test was conducted. *P* < 0.0001 (****).

### Deletion of REV1 enables a more rapid evolution of resistance to caspofungin

The increased rate of *URA3* mutation seen in response to the deletion of *REV1* motivated us to explore whether this increase might affect the rate at which antifungal resistance mutations are acquired, and thus the rate at which antifungal resistance ultimately evolves. We used a laboratory experimental evolution protocol to monitor how quickly the rev1Δ/Δ mutant evolved resistance to a sublethal concentration of the antifungal caspofungin compared to the wild-type strain. Eight isolated colonies for each strain were serially passaged in both the presence or absence of caspofungin, plated on agar plates of rich media containing or lacking caspofungin, and the growth was assessed. We found that the rev1Δ/Δ mutant and wild-type strains grew similarly after passaging in the absence of the drug: passaged rev1Δ/Δ mutant and wild-type strains grew similarly well when plated on rich media, and were similarly unable to grow when plated on media containing caspofungin (**FIGURE 5A; left panel**). When evolved in the presence of caspofungin, both rev1Δ/Δ mutant and wild-type strains grew similarly when plated on rich media, but the rev1Δ/Δ mutant strain grew much more robustly than the wild-type strain when plated on caspofungin-containing media (**FIGURE 5A; right panel**). This trend of the more rapid evolution of resistance to caspofungin was mostly consistent amongst seven of the eight unique passaged isolates of rev1Δ mutants. Further, this more rapid evolution of resistance to caspofungin was reproducible across three distinct experimental evolution assays (**FIGURE S3**). This suggests that the rev1Δ/Δ strain evolved caspofungin resistance more readily than the wild-type strain upon passaging in the presence of this antifungal, and this increase in resistance is consistent with our findings in the 5-FOA assay, which found that the rev1Δ/Δ mutant had an increased rate of mutagenesis.

**Figure 5.**
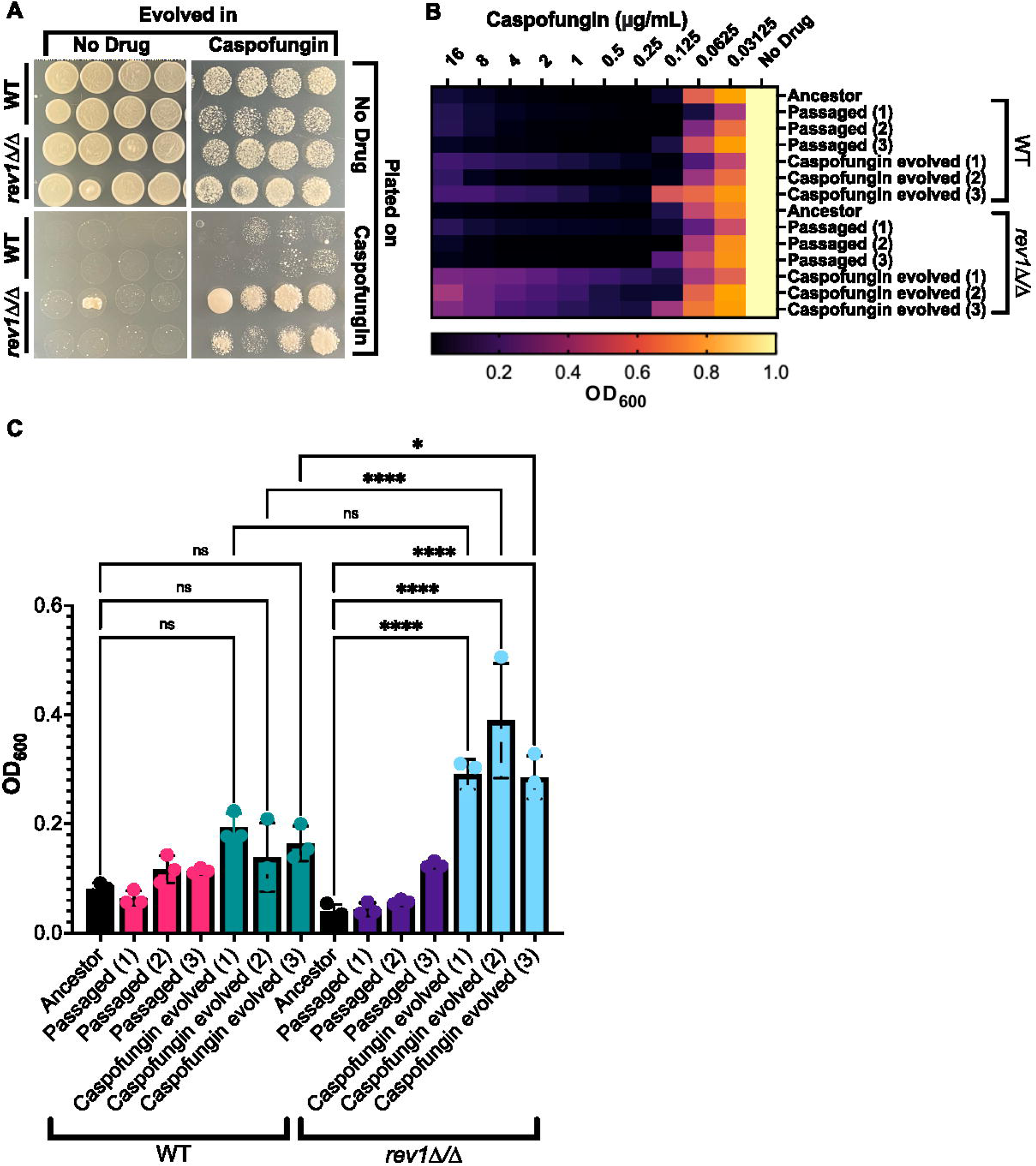
The *rev1*Δ mutant evolves resistance to caspofungin faster than a wild-type strain. **A.** Plate images depict a comparison between the wild-type and *rev1*Δ mutant strains when evolved and plated in the presence and absence of caspofungin. **B**. Heat map depicting evolved wild-type and *rev1*Δ mutant isolates exposed to a concentration gradient of caspofungin. **C**. Optical density of each isolate at the highest concentration of caspofungin. Pairwise comparisons were made between the unevolved and evolved isolates within each strain and between the evolved isolates of both strains. A one-way ANOVA test was conducted. (*) *P* = 0.0494, *P* <0.0001 (****).

Next, we sought to characterize and further quantify the caspofungin resistance seen in the rev1Δ/Δ mutant by performing minimum inhibitory concentration (MIC) assays on individual colonies from these passaged and evolved strains to determine their level of antifungal resistance. We monitored resistance across three single-colony isolates for the passaged strains compared to an unevolved strain. We found that the unevolved isolates of both rev1Δ/Δ and wild-type strains showed similar susceptibility to caspofungin, and the rev1Δ/Δ and wild-type colonies passaged in the absence of the drug also showed similar susceptibility (**FIGURE 5B,C**). When comparing the isolates that were evolved in caspofungin, we found that the rev1Δ/Δ mutant strain colonies were capable of growing at higher concentrations of caspofungin (**FIGURE 5B,C**), suggesting decreased susceptibility to this antifungal across evolved rev1Δ/Δ isolates.

### Deletion of rev1Δ/Δ results in a significant decrease in the abundance of Shm1

Given the unexpected finding that the *C. albicans* rev1Δ/Δ strain more rapidly acquired mutations compared with a wild-type strain, we wanted to assess if this phenotype might be associated with regulatory changes in the rev1Δ/Δ mutant. Thus, we performed proteomics analysis on the rev1Δ/Δ mutant strain compared with the wild-type or *REV1* reconstituted strain in order to gain a more global understanding of what factors may be modulated in response to the deletion of *REV1*. Approximately ∼2,500 proteins were identified in each sample, and only two proteins were found to be significantly modulated in both the comparison between the rev1Δ/Δ mutant and reconstituted strain, and between the rev1Δ/Δ mutant and wild-type strain (**FIGURE 6A, S4, TABLE S1**). Rck2, a putative serine/threonine protein kinase^51^, demonstrated a modest increase in abundance in the rev1Δ/Δ mutant compared to both the reconstituted and wild-type strains and Shm1, a serine hydroxymethyltransferase^52^, had a very clear signature of a significant decrease in abundance (**FIGURE 6A, S4, TABLE S1**).

**Figure 6.**
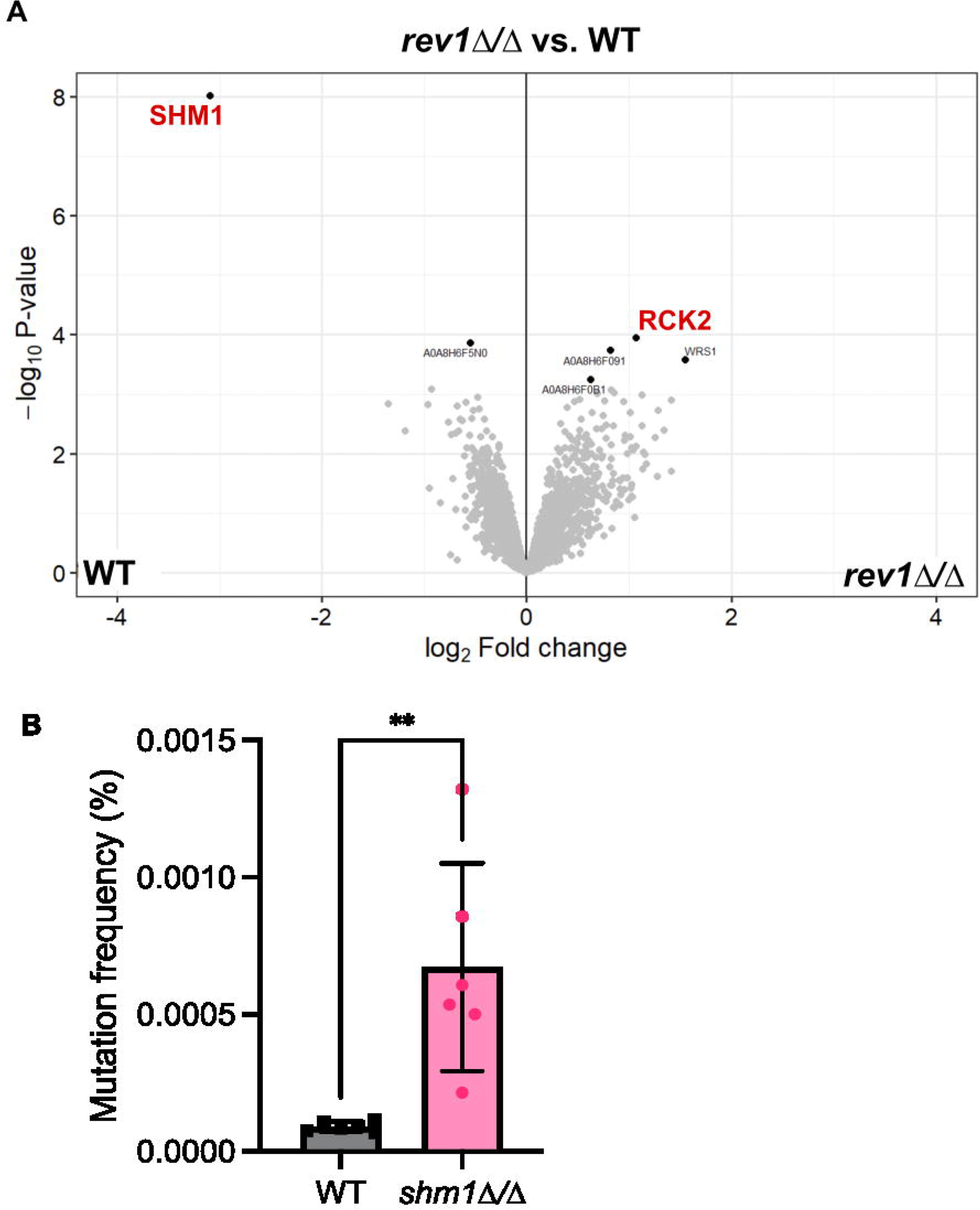
Identification and characterization of Shm1. **A**. Volcano plot of all proteins identified via proteomic analysis of WT, rev1Δ/Δ strains. Red lettering shows the downregulated protein CaJ7.0385 (Shm1) and the upregulated protein Rck2. **B**. Quantified mutation rate based on the number of 5-FOA colonies as a fraction of total number of cells plated. An unpaired t-test was conducted. *P* < 0.005 (**).

### Deletion of SHM1 results in an increased rate of mutation

Based on the finding of Shm1 being significantly depleted in abundance upon deletion of *REV1*, we next sought to characterize Shm1 in *C. albicans*. *SHM1* encodes a predicted serine hydroxymethyltransferase involved in glycine metabolism ^52^, but to date has not been robustly characterized in *C. albicans*. To determine whether *SHM1* is involved in an increased mutation rate, similar to the revΔ/Δ strain, we deleted *SHM1* in *C. albicans* using CRISPR techniques and monitored the mutation rate via 5-FOA assays. Similar to what we observed in the rev1Δ/Δ deletion mutation, deletion of *SHM1* similarly results in an increased mutation rate (**FIGURE 6B**). This suggests that similar to Rev1, Shm1 — itself downregulated upon deletion of *REV1* — also plays a role in the mutation rate of *C. albicans*. In *S. cerevisiae*, *SHM1* has an annotated role in genome stability and chromosome/plasmid maintenance^53^. If this role is conserved in *C. albicans,* this might provide insight into the increase in mutation rate observed in the *C. albicans* shm1Δ/Δ deletion mutation.

## Discussion

Here, we investigate the role of error-prone polymerases in DNA damage and antifungal drug resistance in the fungal pathogen *C. albicans*. We confirm that antifungal agents such as fluconazole and caspofungin can induce DNA double-strand breaks, and find that several predicted error-prone polymerases and related factors were upregulated in these DNA-damaging conditions. Following up on the putative error-prone polymerase *REV1*, we find that deletion of this gene results in sensitivity to DNA damage, and, unexpectedly, increased frequency of mutation, resulting in more rapid evolution of resistance to antifungal drugs. We performed proteomics analysis as a means to assess the mechanism underpinning *REV1*-mediated mutagenesis and drug resistance and identified a previously uncharacterized factor, *SHM1*, which, when deleted in *C. albicans*, similarly results in an increased frequency of mutation. Together, this work describes the importance of previously uncharacterized factors in *C. albicans’* DNA damage response pathway, mutagenesis, and drug resistance.

In bacterial species, error-prone polymerases are known to become upregulated under conditions of DNA damage to prevent cell cycle arrest^20,54^. Here, we find that putative error-prone polymerases are similarly upregulated in *C. albicans* in response to DNA-damaging conditions. While this phenomenon had not previously been investigated in *C. albicans,* previous research in *S. cerevisiae* highlighted the role of *REV1* as an essential component of the translesion DNA synthesis pathway, whose expression is induced in the presence of DNA damage^55^. Further, research in *S. cerevisiae* indicates that the deletion of *REV1* results in increased sensitivity to MMS, UV radiation, and oxidative stress^45,56^. Given its role in error-prone DNA synthesis, the absence of *REV1* in *S. cerevisiae* also confers a decreased mutation rate^57^. In agreement with what was seen in *S. cerevisiae*, we saw a significant sensitivity to treatment with MMS upon deletion of the *REV1* ortholog in *C. albicans*, yet, surprisingly, and in contrast to work in *S. cerevisiae*, we found an increase in mutation rate in the rev1Δ/Δ mutant compared with a wild-type strain. This suggests that *REV1* may have a function in *C. albicans* which differs from that in *S. cerevisiae*, and led us to question what may be occurring in the absence of the gene that might be causing this observed increased mutation rate phenotype.

To understand what was occurring on a cellular systems level upon deletion of *REV1* in *C. albicans,* and to help explain the increased mutation rates and more rapid evolution of caspofungin resistance, we performed proteomic analysis on the *rev1*Δ mutant, *REV1* reconstituted strain, and wild-type strain of *C. albicans*, and found two proteins were significantly modulated in the mutant compared with wild-type or reconstitute strain. Of these two proteins, one, Shm1, was of particular interest as its homolog in *S. cerevisiae* is plays a role in chromosome and plasmid maintenance^53^, and genome stability is known to play an important role antifungal drug resistance^58–64^. Thus the downregulation of the Shm1 protein, putatively involved in chromosome maintenance, may be an example of genomic instability that is induced upon the deletion of *REV1*, leading to an increased mutation rate and more rapid acquisition of antifungal drug resistance.

While Rev1 and other putative error-prone polymerases have remained largely uncharacterized in *C. albicans*, other DNA damage repair factors have well-established roles in the response to antifungal drugs. Previous work in *C. albicans* has demonstrated that deletion of genes involved in double-strand break repair (including *MRE11* and *RAD50*) leads to an increase in genome instability, and increased susceptibility to the antifungal fluconazole^65^. Deletion of *RAD50* and *RAD52* has also been demonstrated to lead to a significant decrease in the fungicidal activity of antifungal agents^31^, and perturbation of factors involved in DNA damage repair in the fungal pathogen *Cryptococcus neoformans* leads to increased susceptibility to certain antifungal agents^66^. Conversely, deletion of genes involved in mismatch repair (*MSH2* and *PMS1*) leads to more rapid acquisition of drug resistance^65^, similar to what we observed upon deletion of *REV1* and *SHM1*. Strains of *Nakaseomyces glabrata* carrying mutations in *MSH2* also exhibit a higher likelihood of breakthrough antifungal treatment *in vitro* and *in vivo*, highlighting the role of mismatch repair in influencing antifungal drug susceptibility ^67^. While our work uncovered an unexpected phenotype of increased frequency of mutations and evolution of drug resistance upon deletion of *REV1* (in *S. cerevisiae,* deletion of the ortholog leads to decreased mutation rate^57^), previous work has shown significant transcriptional rewiring of the DNA damage response between *S. cerevisiae* and the closely-related pathogen *N. glabrata*^68^, suggesting that our findings may reflect similar rewiring in *C. albicans*.

## Materials and Methods

### Strains, media and growth conditions

*C. albicans* cells were grown at 37°C in Yeast Peptone Dextrose (YPD) broth and YPD plates supplemented with 250µg/mL nourseothricin (NAT) for plasmid selection. *Escherichia coli* DH5α cells were grown at 37°C in Lysogeny Broth (LB) and LB plates supplemented with 100mg/mL ampicillin (AMP) and 250µg/mL nourseothricin (NAT) for plasmid selection. The *C. albicans* parental strain used to generate the mutant strains in this study was SN152^69^, and the Gam-GFP *C. albicans* strain was previously described^46^. Strains and plasmids are described in Table S2.

### Strain generation via CRISPR

The deletion strains were generated using the CRISPR-based gene drive platform, as previously described^48,70^. CRISPR gene drive plasmid was constructed using an existing plasmid backbone and a synthesized gene block (Integrated DNA Technologies) specific to the target gene. The existing plasmid backbone and the synthesized gene block were ligated using Gibson Assembly^71^. The ligated plasmid was transformed into DH5α cells and cells were plated on LB plates containing ampicillin for plasmid selection. The plasmid was then purified using the GeneJet Plasmid Miniprep Kit (Thermo Scientific). Following the PCR confirmation of each plasmid, the gene drive plasmid was then digested with PacI and transformed into a diploid *C. albicans* strain using regions of homology upstream and downstream of the target gene, ensuring the complete integration of the gene drive and the deletion of the target gene. Integration of the gene drive in place of each target gene was confirmed via PCR.

### C. albicans transformation

Plasmids were transformed into *C. albicans* via a chemical transformation strategy, as previously described in detail^48,72^. Miniprepped plasmids were linearized via restriction digest mix using the PacI enzyme. A transformation master mix containing 800µl of 50% polyethylene glycol (PEG 3350), 100µl of 10X Tris-EDTA (TE) buffer solution, 100µl of 1M lithium acetate (LiAc), 40µl of 10 mg/mL salmon sperm DNA, and 20µl of 1M dithiothreitol (DTT) was added to *C. albicans* culture and plasmid DNA and incubated at 30°C for 1h, then heat-shocked at 42°C for 45 min. Cells were washed with YPD and grown in fresh YPD for 4 hours. Transformed cells were plated on YPD media containing 250µg/mL NAT, and grown at 30°C for 2 days.

### Microscopy

The *C. albicans* strain containing the *GAM-GFP* construct^46^ was cultured overnight in 5mL YPD shaking at 30LJ. Two overnights were set for each condition tested, and 100µg/mL doxycycline (DOX) was added to half of the cultures. 5mL YPD was included as a ‘no growth’ control. 20mM H_2_O_2_, 0.5µg/mL caspofungin, and 100µg/mL fluconazole were added to the appropriate overnight cultures. The cultures were incubated by shaking at 300C for an additional 4 hours. Following the incubation, 500µL of each culture was transferred to a 1.5mL microcentrifuge tube and centrifuged at 12,000 × g for 10 minutes, after which the supernatant was discarded. The pellet was resuspended in 1mL of 15µg/mL DAPI stain. The resuspended pellets were then incubated in the dark at room temperature for 30 minutes. 2µL of the cell suspension was pipetted onto a microscope slide and visualized via confocal laser scanning microscopy (Leica TCS SP5) at 63x magnification. Leica Application Suite X (LAS X) Life Science Microscope Software Platform was used for image analysis and quantification.

### RNA extraction and reverse transcription quantitative PCR

*C. albicans* strains were cultured overnight in 5mL YPD shaking at 30LJ. OD_600_ of each culture was measured and subcultured in fresh media to a concentration of OD_600_ = 0.05 and grown for approximately 2-3 hours until OD_600_ of 0.2. Once the desired OD_600_ was achieved, cells were then exposed to 0.01% MMS, 0.02% MMS, or 0.5µg/mL caspofungin. The cells were grown in the presence of the respective drugs for 1-2 hours. The cultures were then pelleted and frozen at -80LJ. RNA was extracted using the RNeasy Mini Kit from Qiagen. Samples were then processed at the Advanced Analysis Centre at the University of Guelph for reverse transcription and real-time quantitative PCR. An RNA ScreenTape assay was performed on all the samples using TapeStation 4150 to assess RNA integrity (Agilent Technologies). cDNA was synthesized using 1,000ng of the RNA sample and was performed using a High-Capacity cDNA Reverse-Transcription kit from (Applied BioSystems). This was done by creating a reaction mixture of 10X reverse-transcription buffer, 25X dNTPs at a concentration of 100mM, random primers, MultiScribe Reverse Transcriptase, nuclease-free water, and 1µg of the RNA sample. The reverse transcription reaction conditions were as follows: 10 minutes at 250C, 120 minutes at 37LJ, and 5 minutes at 850C. QuantStudio 7 Pro Real-Time PCR system (Thermo Fisher Scientific) was used to conduct the real-time PCR assays. The reaction mixture was comprised of 2X SsoAdvanced Universal Inhibitor-Tolerant SYBR supermix (Bio-Rad), forward and reverse primers each at 5µM, nuclease-free water, and cDNA. The cycle conditions were as follows: 3 minutes at 980C polymerase activation step, 40 cycles of a two-step qPCR (10 seconds of 980C denaturation, 30 seconds of 600C combined annealing and extension). Expression levels were calculated and assessed according to the comparative CT method^73^. The expression levels of each gene of interest were compared to the housekeeping gene *ACT1*^74^. The ΔCT values in the samples treated with MMS or caspofungin were compared to the ‘no drug’ samples to obtain ΔΔCT and fold change in expression values^73^. All graphs and statistical analysis were performed using GraphPad Prism v.9.4.1.

### MMS serial dilution spot assay

*C. albicans* strains were cultured overnight in 5mL YPD shaking at 300C. OD_600_ of each culture was measured and subcultured in fresh media to a concentration of 0.05 and grown for approximately 2-3 hours until OD_600_ of 0.2. Once the desired OD_600_ was achieved, the culture was divided into two tubes, with 0.05% methyl methanesulfonate (MMS) added to one culture, and no drug added to the second culture. The strains were grown for an additional hour. A serial dilution was performed in 1X Phosphate Buffered Saline (PBS) across six columns in a 96-well plate, including a control column containing only 1X PBS, and plated onto YPD agar. The plates were incubated for 16-24 hours static at 300C. The resulting colonies were then counted.

### 5-FOA fluctuation assay

The desired *C. albicans* strains were cultured overnight in 5mL YPD shaking at 300C. OD_600_ of each culture was measured and the strains were subcultured in fresh media to a concentration of 0.05 and grown for approximately 2-3 hours until OD_600_ of 0.2 and grown for an additional two hours. Following the incubation, the OD_600_ of each culture was measured and the cells were subsequently centrifuged at 5,000 rpm for 10 minutes. The supernatant was discarded and the OD_600_ values were adjusted to the same value in 1X PBS. 100µL of the cell suspension was plated in triplicate for each condition onto 5-fluoroorotic acid (5-FOA) agar plates. The plates were then incubated for 24-48 hours, static at 300C. The resulting colonies were then counted.

### Experimental evolution

Experimental evolution experiments were adapted from previously described protocols^75^. The desired *C. albicans* strains were removed from glycerol stocks stored at -800C, and plated for isolated colonies on YPD agar. 8 isolated colonies for each strain were then cultured overnight in 5mL YPD shaking at 300C. OD_600_ of each culture was measured and OD_600_ values were adjusted to 0.05 in 1mL fresh YPD. 100µL of the diluted cells were added to 900µL casitone, supplemented with 100µg/mL streptomycin in a 96-well deep well plate. Each colony was cultured in duplicate in the 96-well deep well plate, with half being cultured in casitone supplemented with 100µg/mL streptomycin, and the other half being cultured in casitone, streptomycin and 0.25µg/mL caspofungin. Following inoculation, the 96-well plate was covered with a BreatheEasy membrane, wrapped in aluminum foil, and incubated statically at 300C for 7 days. Following the incubation period, 100µL of each culture was removed to create glycerol stocks. 5µL of the evolved culture was plated on a casitone agar plate, and a casitone agar plate supplemented with 0.25µg/mL caspofungin. The plates were grown static at 300C for 16-24 hours and the resulting colonies were analyzed. The remaining culture was centrifuged in the 96-well deep well plate at 1500 rpm for 2 minutes. The supernatant was removed and the resulting pellets were resuspended in fresh casitone, streptomycin, and caspofungin.

### Antifungal minimum inhibitory concentration assays

This protocol was adapted from previously described minimum inhibitory concentration (MIC) assays^58^. The caspofungin MIC assays were performed in flat-bottom 96-well plates. A solution of 32µg/mL caspofungin was prepared in YPD, of which 100µL was added to the first column of the 96-well plate. A two-fold serial dilution was performed across the columns creating a gradient of caspofungin ranging from 16µg/mL to 0µg/mL. The desired evolved isolates were cultured overnight in 5mL YPD shaking at 30°C. OD_600_ of each culture was measured and the cultures were diluted to an OD_600_ of 0.1 in fresh YPD. The diluted culture was mixed into the plates in equal volumes creating a starting cell concentration of OD_600_ = 0.05 for each strain. Following the serial dilution of caspofungin and the addition of the culture, the final volume in each well is 200µL. The plates were then incubated statically at 37°C for 48 hours, after which the optical density (OD_600_) was measured using the Infinite 200 PRO microplate reader (Tecan).

### Proteome extraction

The protocol for protein extraction was adapted from previously described protocols^76^. *C. albicans* strains were cultured overnight in 5mL YPD. The following day, cell pellets were collected, washed and exposed to a protease inhibitor. The cells were then lysed using a probe sonicator and 20% sodium dodecyl sulphate (SDS) was added to the samples for a final concentration of 2% SDS. Next,1µL of 1M dithiothreitol (DTT) per 100µL of the sample was added for a final concentration of 10mM DTT. The samples were then incubated at 95°C for 10 minutes with shaking at 800 rpm. Next, the samples were cooled to room temperature and 1µL iodoacetamide (IAA) was added per 10µL of sample for a final concentration of 55mM IAA. Following a 20-minute incubation at room temperature, 100% ice-cold acetone was added to each sample and samples were stored at -20°C overnight. The following day, the samples were centrifuged to pellet the precipitate, which was then washed in 80% ice-cold acetone. The pellet was then left to air dry. The pellet was re-solubilized in 100µL 8M urea/40mM HEPES. The amount of protein in each sample was then quantified using a Bovine Serum Albumin (BSA) standard curve. Following the protein extraction, the samples were sent to the National Microbiology Laboratory where they further processed the samples, performed mass spectrometry, and completed the proteomic profiling and analysis, as described below.

### Proteomics sample preparation

Each sample (100 μg) was brought to 5% Sodium Dodecyl Sulfate (MP Biochemicals) in a total volume of 100 μl. The samples were processed using on-filter digestion with S-Trap mini spin columns (Protifi) according to the manufacturer’s instructions. Chemicals used for filtering on S-Trap columns were Tris(2-carboxyethyl)phosphine, Methyl methanethiosulfonate, Triethylammonium bicarbonate, MgCl2, (Thermo Scientific), Phosphoric acid, and acetonitrile (Fisher Chemicals), Formic acid (Merek), Benzonase (EMD Millipore/Novagen). The samples were digested with Trypsin (Pierce). The digested samples were concentrated to near-dryness (1-5 μl) using a vacuum centrifuge then were labeled with TMT 10plex Isobaric Label Reagent Set (Thermo Fisher Scientific) as per the manufacturer’s protocol, and TMT-labelled samples were mixed one-to-one.

TMT samples mixtures were resuspended in 40 µL of buffer A (20mM ammonium formate,) and fractionated by high-pH C18 reversed-phased liquid chromatography using an Agilent 1260 Infinity liquid chromatography system (Agilent Technologies) equipped with a XBridge C18 3.5-μm, 2.1x100-mm column (Waters) using a linear gradient of 20-56% buffer B (20mM ammonium formate, 90% acetonitrile) over 52.5 min. The twelve TMT fractions were concentrated to near-dryness using vacuum centrifugation, and resuspended in 80 µL of nano buffer A (2% acetonitrile, 0.1% formic acid).

### Proteomics data acquisition

Each TMT fraction was analyzed using a nano-flow Easy nLC 1200 connected in-line to an Orbitrap Fusion Lumos mass spectrometer with a nanoelectrospray ion source at 2.3 kV and a FAIMS Pro module (Thermo Fisher Scientific). The TMT-labeled peptide samples were loaded (2 µl) onto a C18-reversed phase Easy Spray column (50 cm long, 75 µm inner diameter, 2 µm particles; Thermo Fisher Scientific) with 100% nano buffer A. Peptides were eluted using a linear gradient of 3-27% nano buffer B (80% acetonitrile, 0.1% formic acid) over 100 min. Total LC-MS/MS run-time was approximately 150 minutes, including the loading, linear gradient, column wash, and column equilibration.

Data-dependent acquisition was used with three different CV settings on the FAIMS Pro (-40, -55, and -75) with each CV setting taking 1 second for a total of a 3 second cycle time. For each CV, precursor ions were selected and isolated in the quadrupole (0.7 m/z isolation width) and fragmented by CID (35% normalized collision energy), as many as possible within a 1-second interval. MS3 scans were acquired using 10 SPS precursors fragmented by HCD (65% normalized collision energy). The survey scans were acquired in the Orbitrap over a range of m/z 400-1600 with a target resolution of 120,000 at m/z 200, fragment ion (MS2) scans were acquired in the iontrap at the Turbo scan rate, and MS3 scans were acquired in the Orbitrap with a target resolution of 50,000 at m/z 200.

### Proteomics data processing

Raw mass spectrometry data was searched using the Sequest search engine within Proteome Discoverer version 2.5 (Thermo Scientific) against a *C. albicans* (NCBI taxonomy ID: 5476) protein fasta database downloaded from Uniprot. Search results were exported in tab-delimited format for further data processing using the R programming language. All data processing including differential expression analysis was performed using the Bioconductor package DEP^77^. Variance stabilizing normalization was performed as per the vsn packagè, then data was imputed assuming missing not at random (MNAR) by taking random draws from a Gaussian distribution centered around a minimal value. Volcano plots were generated after differential expression analysis using the plot_volcanò function within the DEP package.

## Acknowledgments

We thank Shapiro lab members, past and present, for helpful discussions and support of this project, as well as Dr. Meleah Hickman, Dr. Ognenka Avramovska, and Dr. Aleeza Gerstein for helpful conversations, suggestions, and feedback. We would like to thank Stuart McCorrister in the Mass Spectrometry & Proteomics Core at the National Microbiology Laboratory, Public Health Agency of Canada (PHAC), for acquiring the proteomics data by LC-MS/MS and Sierra Rosiana-Easlick at PHAC for coordination support. We thank the staff at the University of Guelph Advanced Analysis Centre for support with qRT-PCR and microscopy. This work was supported by an NSERC Discovery Grant (RGPIN-2018-4914), an Ontario Early Research Award, and a CIFAR Azrieli Global Scholar award to RSS, and a Genome Canada and Genome Quebec grant (6569) and NSERC Discovery Grant (RGPIN-2020-04844) to CRL. M.R.A-T. was supported by an EvoFunPath Fellow (NSERC CREATE).

---

**Figure S1.**
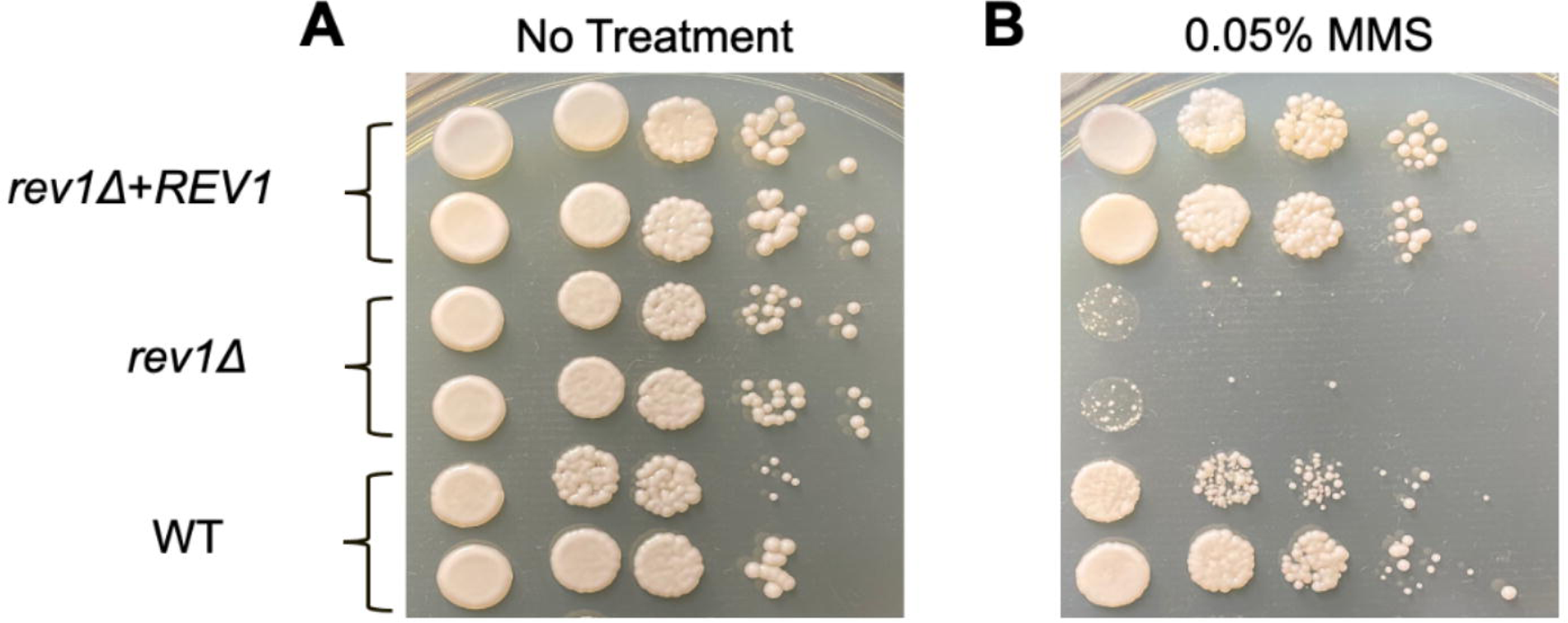
Reconstitution of *REV1* restores wild-type phenotype. (**A and B**). The *REV1* reconstituted strain was grown in the presence of MMS alongside the mutant and wild-type strains and displayed a lack of sensitivity to the DNA-damaging agent.

**Figure S2.**
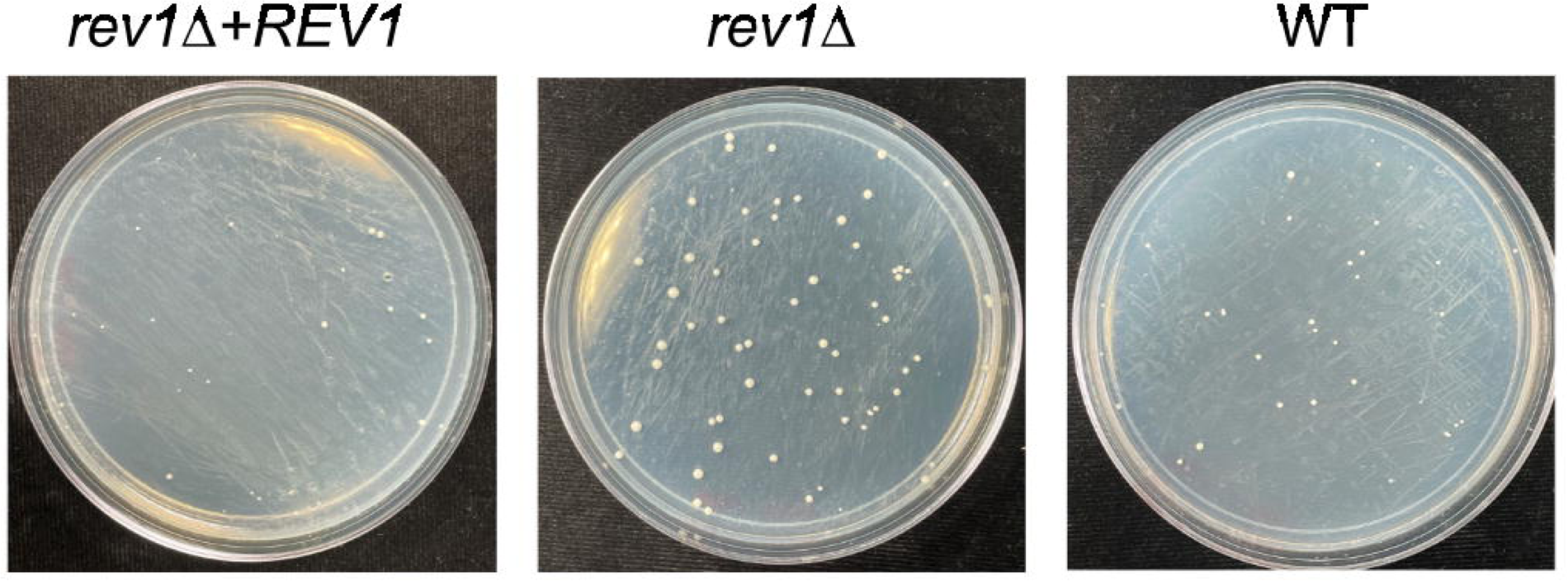
Restoration of *REV1* reverses mutagenic phenotype. Plate images depict *REV1* reconstituted strain, *rev1*Δ mutant, and wild-type strain plated on 5-FOA.

**Figure S3.**
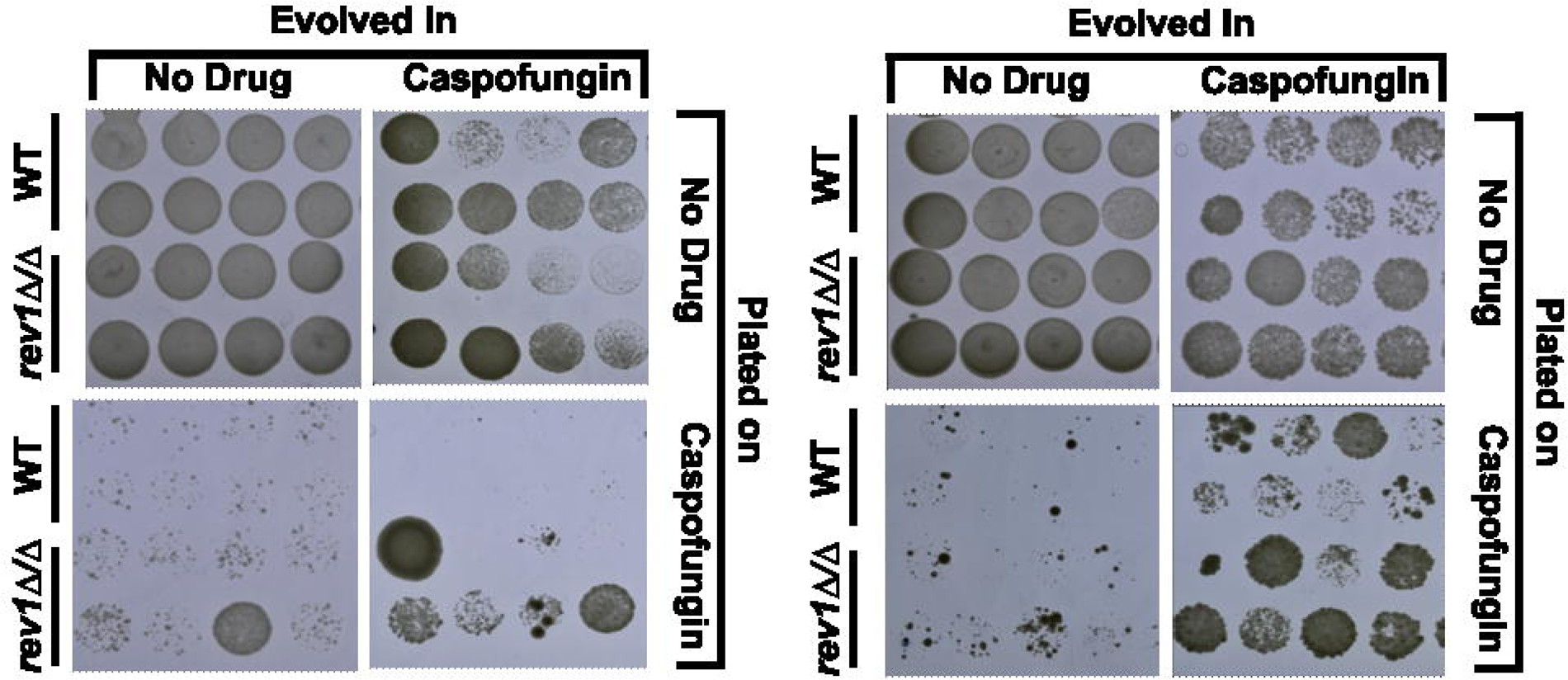
The *rev1*Δ mutant evolves resistance to caspofungin faster than a wild-type strain. **A.** Plate images depict a comparison between the wild-type and *rev1*Δ mutant strains when evolved and plated in the presence and absence of caspofungin, across two additional independent replicates.

**Figure S4.**
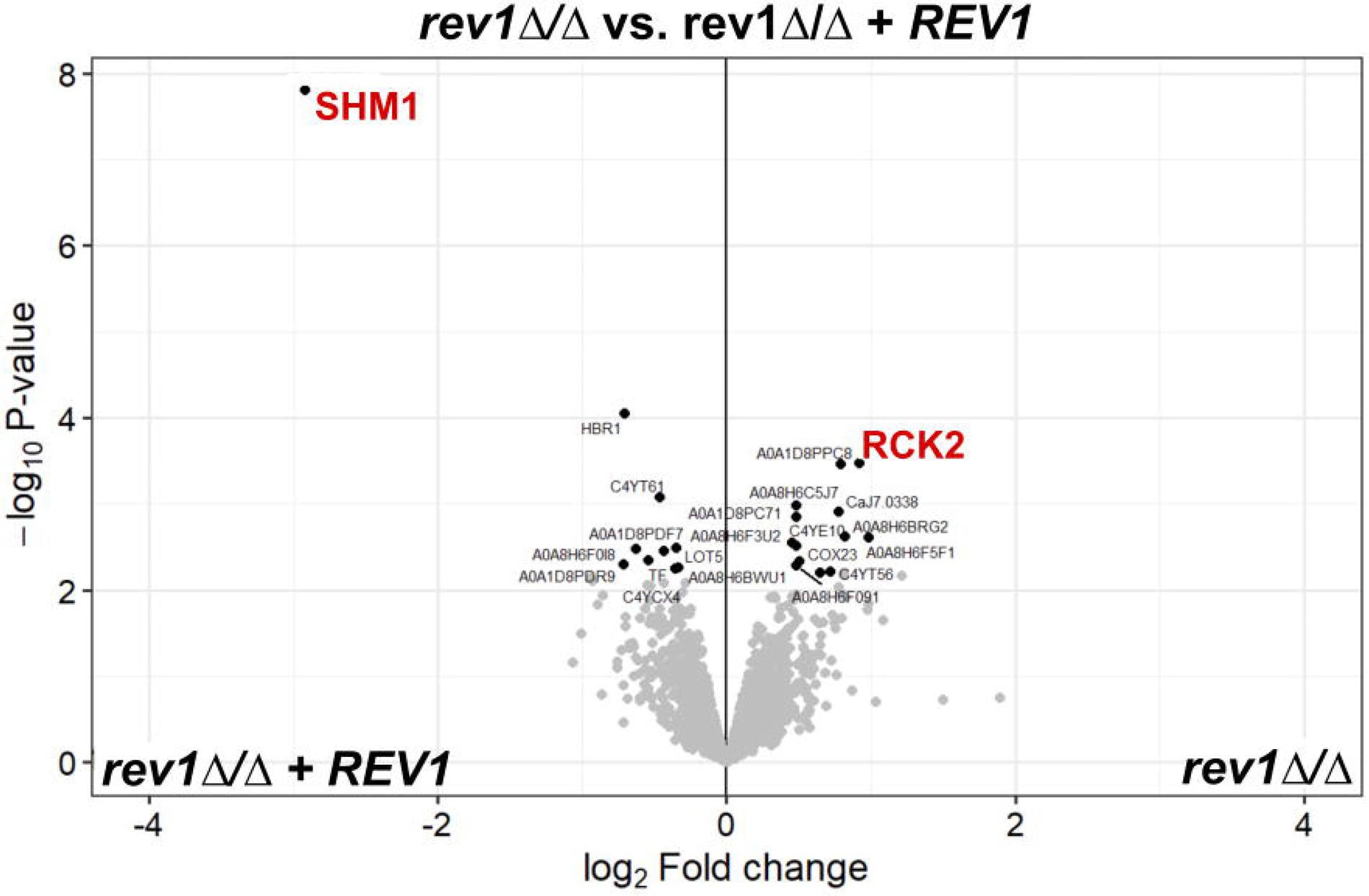
Two proteins were found to be significantly modulated upon deletion of *REV1*. (**A**) Volcano plot of all proteins identified via proteomic analysis of rev1Δ/Δ and *REV1* constituted strains. Red lettering shows the downregulated protein CaJ7.0385 (Shm1) and the upregulated protein Rck2.

## Notes

### Competing Interest Statement

The authors have declared no competing interest.

## References

1. Fisher, M. C. et al. Threats Posed by the Fungal Kingdom to Humans, Wildlife, and Agriculture. MBio 11, (2020).

2. Vallabhaneni, S., Mody, R. K., Walker, T. & Chiller, T. The Global Burden of Fungal Diseases. Infect. Dis. Clin. North Am. 30, 1–11 (2016).

3. Nnadi, N. E. & Carter, D. A. Climate change and the emergence of fungal pathogens. PLoS Pathog. 17, e1009503 (2021).

4. Casadevall, A. Climate change brings the specter of new infectious diseases. J. Clin. Invest. 130, 553–555 (2020).

5. Fisher, M. C. et al. Emerging fungal threats to animal, plant and ecosystem health. Nature 484, 186–194 (2012).

6. Bongomin, F., Gago, S., Oladele, R. & Denning, D. Global and Multi-National Prevalence of Fungal Diseases—Estimate Precision. Journal of Fungi 3, 57 (2017).

7. Pfaller, M. A. & Diekema, D. J. Epidemiology of invasive candidiasis: a persistent public health problem. Clin. Microbiol. Rev. 20, 133–163 (2007).

8. Taei, M., Chadeganipour, M. & Mohammadi, R. An alarming rise of non-albicans Candida species and uncommon yeasts in the clinical samples; a combination of various molecular techniques for identification of etiologic agents. BMC Res. Notes 12, 779 (2019).

9. Miceli, M. H., Díaz, J. A. & Lee, S. A. Emerging opportunistic yeast infections. Lancet Infect. Dis. 11, 142–151 (2011).

10. Pfaller, M. A., Diekema, D. J., Turnidge, J. D., Castanheira, M. & Jones, R. N. Twenty Years of the SENTRY Antifungal Surveillance Program: Results for Candida Species From 1997–2016. Open Forum Infect Dis 6, S79–S94 (2019).

11. Geddes-McAlister, J. & Shapiro, R. S. New pathogens, new tricks: emerging, drug-resistant fungal pathogens and future prospects for antifungal therapeutics. Ann. N. Y. Acad. Sci. 1435, 57–78 (2019).

12. Perfect, J. R. The antifungal pipeline: a reality check. Nat. Rev. Drug Discov. 16, 603–616 (2017).

13. Pappas, P. G. et al. Clinical Practice Guideline for the Management of Candidiasis: 2016 Update by the Infectious Diseases Society of America. Clin. Infect. Dis. 62, e1–50 (2016).

14. Denning, D. W. Echinocandin antifungal drugs. Lancet 362, 1142–1151 (2003).

15. Alexander, B. D. et al. Increasing echinocandin resistance in Candida glabrata: clinical failure correlates with presence of FKS mutations and elevated minimum inhibitory concentrations. Clin. Infect. Dis. 56, 1724–1732 (2013).

16. Kohanski, M. A., Depristo, M. A. & Collins, J. J. Sublethal Antibiotic Treatment Leads to Multidrug Resistance via Radical-Induced Mutagenesis. Mol. Cell 37, 311–320 (2010).

17. Pribis, J. P., Zhai, Y., Hastings, P. J. & Rosenberg, S. M. Stress-Induced Mutagenesis, Gambler Cells, and Stealth Targeting Antibiotic-Induced Evolution. MBio 13, e0107422 (2022).

18. Gutierrez, A. et al. β-Lactam antibiotics promote bacterial mutagenesis via an RpoS-mediated reduction in replication fidelity. Nat. Commun. 4, 1610 (2013).

19. Shapiro, R. S. Antimicrobial-induced DNA damage and genomic instability in microbial pathogens. PLoS Pathog. 11, e1004678 (2015).

20. Long, H. et al. Antibiotic treatment enhances the genome-wide mutation rate of target cells. Proc. Natl. Acad. Sci. U. S. A. 113, 2498–2505 (2016).

21. Bush, N. G., Diez-Santos, I., Abbott, L. R. & Maxwell, A. Quinolones: Mechanism, Lethality and Their Contributions to Antibiotic Resistance. Molecules 25, (2020).

22. Hawkey, P. M. Mechanisms of quinolone action and microbial response. J. Antimicrob. Chemother. 51 **Suppl 1**, 29–35 (2003).

23. Dwyer, D. J., Collins, J. J. & Walker, G. C. Unraveling the physiological complexities of antibiotic lethality. Annu. Rev. Pharmacol. Toxicol. 55, 313–332 (2015).

24. Zhao, X. & Drlica, K. Reactive oxygen species and the bacterial response to lethal stress. Curr. Opin. Microbiol. 21, 1–6 (2014).

25. Foti, J. J., Devadoss, B., Winkler, J. A., Collins, J. J. & Walker, G. C. Oxidation of the guanine nucleotide pool underlies cell death by bactericidal antibiotics. Science 336, 315– 319 (2012).

26. Pérez-Capilla, T. et al. SOS-independent induction of dinB transcription by beta-lactam-mediated inhibition of cell wall synthesis in Escherichia coli. J. Bacteriol. 187, 1515–1518 (2005).

27. Ysern, P. et al. Induction of SOS genes in Escherichia coli and mutagenesis in Salmonella typhimurium by fluoroquinolones. Mutagenesis 5, 63–66 (1990).

28. Thi, T. D. et al. Effect of recA inactivation on mutagenesis of Escherichia coli exposed to sublethal concentrations of antimicrobials. J. Antimicrob. Chemother. 66, 531–538 (2011).

29. Baharoglu, Z. & Mazel, D. Vibrio cholerae triggers SOS and mutagenesis in response to a wide range of antibiotics: a route towards multiresistance. Antimicrob. Agents Chemother. 55, 2438–2441 (2011).

30. Cirz, R. T. et al. Inhibition of mutation and combating the evolution of antibiotic resistance. PLoS Biol. 3, e176 (2005).

31. Belenky, P., Camacho, D. & Collins, J. J. Fungicidal drugs induce a common oxidative-damage cellular death pathway. Cell Rep. 3, 350–358 (2013).

32. Mesa-Arango, A. C. et al. The production of reactive oxygen species is a universal action mechanism of Amphotericin B against pathogenic yeasts and contributes to the fungicidal effect of this drug. Antimicrob. Agents Chemother. 58, 6627–6638 (2014).

33. Hao, B., Cheng, S., Clancy, C. J. & Nguyen, M. H. Caspofungin kills Candida albicans by causing both cellular apoptosis and necrosis. Antimicrob. Agents Chemother. 57, 326–332 (2013).

34. Avramovska, O. & Hickman, M. A. The Magnitude of Candida albicans Stress-Induced Genome Instability Results from an Interaction Between Ploidy and Antifungal Drugs. G3 9, 4019–4027 (2019).

35. Forche, A. et al. Stress alters rates and types of loss of heterozygosity in Candida albicans. MBio 2, (2011).

36. Krutiakov, V. M. Eukaryotic error prone DNA polymerases: suggested roles in replication, repair and mutagenesis. Mol. Biol. 40, 3–11 (2006).

37. Waters, L. S. et al. Eukaryotic translesion polymerases and their roles and regulation in DNA damage tolerance. Microbiol. Mol. Biol. Rev. 73, 134–154 (2009).

38. Kunkel, T. A. DNA replication fidelity. J. Biol. Chem. 279, 16895–16898 (2004).

39. Lawrence, C. W. Cellular roles of DNA polymerase zeta and Rev1 protein. DNA Repair 1, 425–435 (2002).

40. Prakash, S. & Prakash, L. Translesion DNA synthesis in eukaryotes: a one- or two-polymerase affair. Genes Dev. 16, 1872–1883 (2002).

41. Shcherbakova, P. V. & Fijalkowska, I. J. Translesion synthesis DNA polymerases and control of genome stability. Front. Biosci. 11, 2496–2517 (2006).

42. Zhong, X. et al. The fidelity of DNA synthesis by yeast DNA polymerase zeta alone and with accessory proteins. Nucleic Acids Res. 34, 4731–4742 (2006).

43. Larimer, F. W., Perry, J. R. & Hardigree, A. A. The REV1 gene of Saccharomyces cerevisiae: isolation, sequence, and functional analysis. J. Bacteriol. 171, 230–237 (1989).

44. Lemontt, J. F. Mutants of yeast defective in mutation induced by ultraviolet light. Genetics 68, 21–33 (1971).

45. Yao, R., Zhou, P., Wu, C., Liu, L. & Wu, J. Sml1 Inhibits the DNA Repair Activity of Rev1 in Saccharomyces cerevisiae during Oxidative Stress. Appl. Environ. Microbiol. 86, (2020).

46. Thomson, G. J. et al. MetabolismLJinduced oxidative stress and DNA damage selectively trigger genome instability in polyploid fungal cells. The EMBO Journal vol. 38 Preprint at 10.15252/embj.2019101597 (2019).

47. Belenky, P. et al. Bactericidal Antibiotics Induce Toxic Metabolic Perturbations that Lead to Cellular Damage. Cell Rep. 13, 968–980 (2015).

48. Halder, V., Porter, C. B. M., Chavez, A. & Shapiro, R. S. Design, execution, and analysis of CRISPR–Cas9-based deletions and genetic interaction networks in the fungal pathogen Candida albicans. Nat. Protoc. 14, 955–975 (2019).

49. Lang, G. I. & Murray, A. W. Estimating the per-base-pair mutation rate in the yeast Saccharomyces cerevisiae. Genetics 178, 67–82 (2008).

50. Auerbach, P. A. & Demple, B. Roles of Rev1, Pol zeta, Pol32 and Pol eta in the bypass of chromosomal abasic sites in Saccharomyces cerevisiae. Mutagenesis 25, 63–69 (2010).

51. Li, X. et al. The MAP kinase-activated protein kinase Rck2p plays a role in rapamycin sensitivity in Saccharomyces cerevisiae and Candida albicans. FEMS Yeast Res. 8, 715– 724 (2008).

52. McNeil, J. B. et al. Glycine metabolism in Candida albicans: characterization of the serine hydroxymethyltransferase (SHM1, SHM2) and threonine aldolase (GLY1) genes. Yeast 16, 167–175 (2000).

53. Choy, J. S. et al. Genome-wide haploinsufficiency screen reveals a novel role for γ-TuSC in spindle organization and genome stability. Mol. Biol. Cell 24, 2753–2763 (2013).

54. Žgur-Bertok, D. DNA damage repair and bacterial pathogens. PLoS Pathog. 9, e1003711 (2013).

55. Sobolewska, A., Halas, A., Plachta, M., McIntyre, J. & Sledziewska-Gojska, E. Regulation of the abundance of Y-family polymerases in the cell cycle of budding yeast in response to DNA damage. Curr. Genet. 66, 749–763 (2020).

56. Svensson, J. P. et al. Genomic phenotyping of the essential and non-essential yeast genome detects novel pathways for alkylation resistance. BMC Syst. Biol. 5, 157 (2011).

57. McDonald, M. J. et al. Mutation at a distance caused by homopolymeric guanine repeats in Saccharomyces cerevisiae. Sci Adv 2, e1501033 (2016).

58. Gervais, N. C. et al. Development and applications of a CRISPR activation system for facile genetic overexpression in Candida albicans. G3 (2022) doi:10.1093/g3journal/jkac301.

59. Todd, R. T., Wikoff, T. D., Forche, A. & Selmecki, A. Genome plasticity in Candida albicans is driven by long repeat sequences. Elife 8, (2019).

60. Forche, A. et al. Rapid Phenotypic and Genotypic Diversification After Exposure to the Oral Host Niche in Candida albicans. Genetics 209, 725–741 (2018).

61. Yang, F. et al. The fitness costs and benefits of trisomy of each Candida albicans chromosome. Genetics 218, (2021).

62. Selmecki, A., Forche, A. & Berman, J. Aneuploidy and isochromosome formation in drug-resistant Candida albicans. Science 313, 367–370 (2006).

63. Todd, R. T. & Selmecki, A. Expandable and reversible copy number amplification drives rapid adaptation to antifungal drugs. Elife 9, (2020).

64. Kwon-Chung, K. J. & Chang, Y. C. Aneuploidy and drug resistance in pathogenic fungi. PLoS Pathog. 8, e1003022 (2012).

65. Legrand, M., Chan, C. L., Jauert, P. A. & Kirkpatrick, D. T. Role of DNA mismatch repair and double-strand break repair in genome stability and antifungal drug resistance in Candida albicans. Eukaryot. Cell 6, 2194–2205 (2007).

66. Jung, K.-W. et al. Rad53- and Chk1-Dependent DNA Damage Response Pathways Cooperatively Promote Fungal Pathogenesis and Modulate Antifungal Drug Susceptibility. MBio 10, (2019).

67. Healey, K. R. et al. Prevalent mutator genotype identified in fungal pathogen Candida glabrata promotes multi-drug resistance. Nat. Commun. 7, 11128 (2016).

68. Shor, E., Garcia-Rubio, R., DeGregorio, L. & Perlin, D. S. A Noncanonical DNA Damage Checkpoint Response in a Major Fungal Pathogen. MBio 11, (2020).

69. Homann, O. R., Dea, J., Noble, S. M. & Johnson, A. D. A phenotypic profile of the Candida albicans regulatory network. PLoS Genet. 5, e1000783 (2009).

70. Shapiro, R. S. et al. A CRISPR–Cas9-based gene drive platform for genetic interaction analysis in Candida albicans. Nature Microbiology 3, 73–82 (2018).

71. Gibson, D. G. et al. Enzymatic assembly of DNA molecules up to several hundred kilobases. Nat. Methods 6, 343–345 (2009).

72. Wensing, L. & Shapiro, R. S. Design and Generation of a CRISPR Interference System for Genetic Repression and Essential Gene Analysis in the Fungal Pathogen Candida albicans. Methods Mol. Biol. 2377, 69–88 (2022).

73. Schmittgen, T. D. & Livak, K. J. Analyzing real-time PCR data by the comparative C(T) method. Nat. Protoc. 3, 1101–1108 (2008).

74. Nailis, H., Coenye, T., Van Nieuwerburgh, F., Deforce, D. & Nelis, H. J. Development and evaluation of different normalization strategies for gene expression studies in Candida albicans biofilms by real-time PCR. BMC Mol. Biol. 7, 25 (2006).

75. Avramovska, O., Smith, A. C., Rego, E. & Hickman, M. A. Tetraploidy accelerates adaptation under drug selection in a fungal pathogen. Frontiers in Fungal Biology 3, (2022).

76. Ball, B. & Geddes-McAlister, J. Quantitative Proteomic Profiling of Cryptococcus neoformans. Curr. Protoc. Microbiol. 55, e94 (2019).

77. Zhang, X. et al. Proteome-wide identification of ubiquitin interactions using UbIA-MS. Nat. Protoc. 13, 530–550 (2018).

